# MATRIX: Rapid Quantification of Total and Active Microbial Cells with Single-Cell Phenotypes for Environmental Microbiomes

**DOI:** 10.64898/2026.03.16.712149

**Authors:** Milena Gonzalo, Xipeng Liu, Yann S. Dufour, Ashley Shade

## Abstract

Quantifying the abundance and activity of bacteria within populations and communities is fundamental to systems microbiology and microbiome research. Yet direct microscopic cell counting remains low-throughput, labor-intensive, and prone to user variability, leading many researchers to rely on indirect proxies such as optical density or multicopy marker-gene quantification. These indirect approaches do not distinguish between active and inactive cells and can obscure ecological interpretation. Here, we introduce MATRIX (Microbial Activity and Total cell quantification via Rapid Imaging and eXtraction), an efficient workflow that integrates sample extraction, fluorescence staining, automated microscopy and image analysis, and Bayesian statistical inference to quantify total and redox-active cells and derive single-cell measurements for environmental microbial populations and communities. We demonstrate its reproducibility and versatility using both cultured isolates and high-diversity soil communities. The resulting quantitative, phenotypic datasets provide rapid, direct measurements of population of community size and activity, enabling well-powered analyses that strengthen mechanistic insight into microbial responses and improve the ecological grounding of microbiome studies.

**Importance:** Microbiome studies commonly rely on relative abundance data, which cannot distinguish whether compositional shifts reflect true population growth, declines in total community size, or both. Without explicit measurements of population and community sizes, mechanistic interpretation of microbiome dynamics remains incomplete. Here we present a rapid, throughput workflow, MATRIX, that quantifies both total and redox-active bacterial cells from environmental samples. By integrating single-cell phenotypes with community-level metrics, this approach anchors microbiome datasets in direct ecological accounting rather than proxies. These measurements can clarify whether observed changes in community structure represent shifts in abundance, activity, or both, improving inference about microbial responses to stress or environmental change. MATRIX therefore offers an efficient way to incorporate quantitative ecology into systems-microbiology and microbiome studies and to strengthen the link between microbial cellular physiology, community dynamics, and eco-system function.

**Graphical Abstract:** 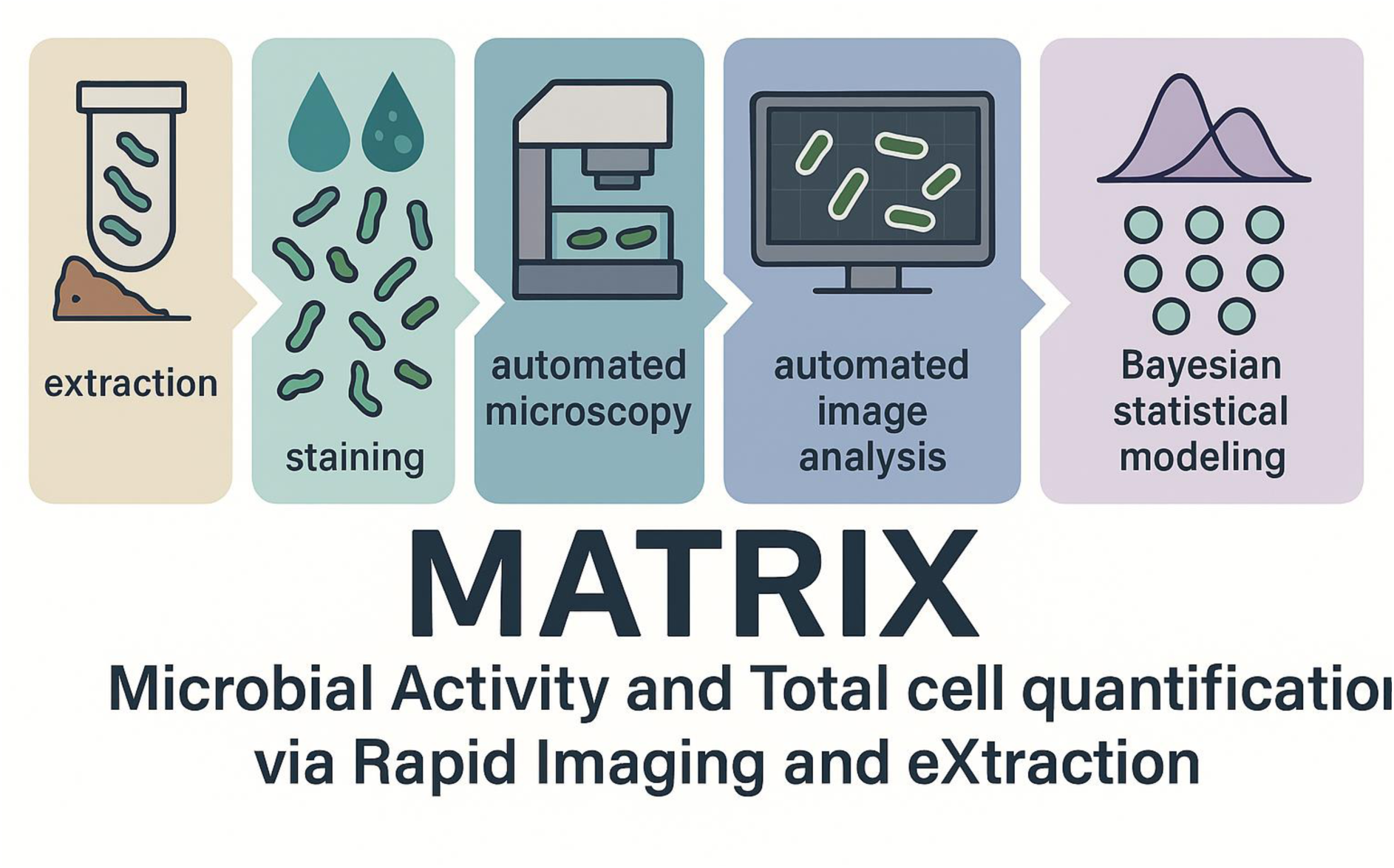

## Introduction

A critical observed variable in systems microbiome science, including microbial ecology, is population or community size (1). This variable quantifies the number of individual cells that comprise a population or consortium living in a particular habitat at a particular moment. Changes in the sizes of microbial populations or communities across space, time, or in response to perturbations, reflect outcomes of key ecological processes, such as deterministic or stochastic dynamics attributable to growth and death (2). Understanding how community size responds to changing host or environmental conditions is essential for interpreting shifts in community structure and function, including those inferred from ‘omics profiles. An increase in a taxon’s relative abundance may reflect its own growth or the loss of other taxa. Thus, relative abundance can change even when a taxon’s absolute abundance remains constant, and the interpretation is muddied without additional information about community and population sizes.

The cultivation-independent assessment of microbiome structure and function has become largely automated high-throughput. However, the direct quantification of community size, sometimes called quantitative microbiome profiling (3, 4), can be tedious, and is therefore less frequently performed. Indirect estimates of community size, such as quantitative PCR of a multicopy marker gene (e.g., 16S rRNA gene, 18S rRNA gene, or intergenic spacer genes, e.g., (5)) or spike-ins of nucleic acids of known concentrations are commonly used (as recently discussed in (6)). Yet these approaches can present challenges when correcting or standardizing compositional microbiome data. First, they are not experimentally independent from the ‘omics workflows they aim to normalize, which can amplify errors and biases (7). Second, multicopy marker genes add additional, often uncorrectable biases due to variation in copy numbers across lineages, making interpretation difficult when relative compositions shift alongside changes in population size (8–10). Third, these methods quantify nucleic acids rather than individual cell counts; cells differ in genome size and nucleic acid content depending on cell sizes (11), taxonomic identity (12) and growth rate at the time of sampling (summarized in (13)). While quantitative approaches have been shown to improve accuracy for comparative microbiome profiling (e.g., (14)), indirect methods remain common due to their ease of use, cost, and compatibility with existing sequencing pipelines.

Another major challenge when studying microbiomes is the ability to discriminate and quantify active members. Microbiome assessment based on DNA markers include signal from dead and inactive members, which can contribute substantially to community measurement (as summarized in (15, 16)). Thus, many of the members and genes detected by marker gene and metagenome sequencing may not be functioning *in situ*. Even activity-referenced microbiome assessment, like metatranscriptomics or proteomics, do not directly quantify the total or active number of organisms (16). As a result, it is often unclear whether compositional shifts in these ‘omics-based functional profiles co-occur with cell death (negative selection) or growth (positive selection). Distinguishing these opposing selection mechanisms can provide insight into ecological consequences such as niche availability, inter-microbial competition, and host interactions.

To discriminate and quantify active versus inactive microbial cells, fluorescence-based assays have long been used (17). These assays stain the cells with a fluorescence signal proportional to their redox activity, and individual cell measurements are captured either by flow cytometry or direct microscopy. However, these protocols are laborious and time consuming, and flow cytometry requires access to specialized instrumentation. As a result, even though these methods provide more quantitative information, they are less favored when compared to high-throughput of ‘omics workflows that can rapidly generate large microbiome datasets.

To address the need for improved access to quantitative microbiome data, we present a rapid, largely automated workflow, MATRIX (Microbial Activity and Total cell quantification via Rapid Imaging and eXtraction), for enumerating total and active microbial cells in mixed environmental communities, with a focus on bacteria. In addition to total and active cell counts, the single-cell image analysis approach provides morphological and fluorescence-intensity information that offers additional biological context for interpreting how cells physiologically respond to environmental changes. The automation allows for the generation of large, reproducible, and quantitative microbial population and community datasets. These data permit well-powered experimental designs and provide an independent, direct assessment of microbiome population or community size, which can then be used to interpret with greater precision the patterns observed in ‘omics-based microbiome diversity.

## Materials and Methods

### Bacterial cell staining

Two types of dyes were used to stain live cells for direct counting and assessment of redox activity. SYBR^TM^ Green I (Invitrogen, CA, USA), which stains all DNA-containing cells (18), was used to quantify total cell counts. Redox Sensor Green (RSG, BacLight RedoxSensor green vitality kit, Invitrogen), which is reduced into its fluorescent form by bacterial reductases (19), was used to quantify respiration-active cell counts and their relative redox potential. Briefly, cells were mixed with either 1 μM of SYBR or 1 μM of RSG and then incubated at room temperature in the dark for 15 minutes according to manufacturer’s instructions.

### Automated microscopy

Automated microscopy was performed using the QUANTOM Tx ^TM^ microbial cell counter (Q100001, Logos Biosystems Inc., South Korea). The following protocols were adapted from the Logos Biosystems User Manual (20) that advises on the rapid quantification of pure bacterial cultures in clinical diagnostic settings. Ten microliters of stained cells were mixed with 10 μL of 100% glycerol and 6 μL of this mixture was loaded into a counting slide (Q12001), centrifuged at 650 x *g* for 10 min in the QUANTOM^TM^ slide Centrifuge (Q10002) to immobilize and uniformly distribute the cells on the counting slide. The slides were loaded into the microbial cell counter, which captures fluorescence signal with an excitation of 470/30 nm and an emission of 530/50 nm, with an excitation LED source set at level of 5. Manufacturer’s recommendations were followed to ensure a concentration between 2 × 10^5^ and 1×10^9^ cells/mL, a loading volume of 6 μL, and a maximum centrifuge 650 x *g*.

### Image analysis

For each assay, ten discrete images are captured in succession per slide. Each image was visually inspected for appropriate concentration, sufficient dispersion of cells, and detection of apparent aggregations or artifacts. We developed the MATRIX image analysis step to address the specific challenges in detecting fluorescence signal from individual cells in our environmental samples and quantify parameters of interests. The background of each image was flattened, centered at zero, and normalized to determine a threshold above 6 standard deviations for fluorescent signal detection. Objects close to each other were segmented using the watershed algorithm. After filtering high frequency noise, coordinates, morphological parameters, and fluorescence intensity were calculated for each object. Objects intersecting the image borders, containing saturated pixels, or out of focus were flagged for further inspections.

For each detected SYBR object, area, long and short axis lengths, eccentricity, and solidity. The volume of each SYBR object was estimated by assuming rotational symmetry along the major length axis. For each SYBR and RSG object, we also detected fluorescence (total per object). In addition, several parameters were calculated per image: the total number of cells was determined by the SYBR counts, and the total number of active cells was determined from the RSG stained cells. In addition, several parameters were calculated per sample (here, soil or culture): the concentration of total cells from the SYBR counts, the concentration of active cells from the RSG counts, the proportion of active cells, and the total fluorescence intensity.

### Culture-based assays

To first validate the automated cell imaging and counting method with pure cultures of bacteria, we assessed changes in active and total cell counts and distribution of activity intensities over the growth curves of commercially available bacterial strains. *Escherichia coli* and *Pseudomonas protegens* were purchased from the Belgian Coordinated Collections of Microorganisms Laboratory of Microbiology, Department of Biochemistry and Microbiology, Faculty of Sciences of Ghent University (BCCM/LMG) bacterial collection. Each strain was inoculated from glycerol stock into Luria-Bertani (LB) agar and grown at 28 °C for 48 hours. One colony was transferred to 50 % liquid LB and grown for 24 hours at 28 °C with shaking at 200 rpm. Optical density at 600 nm (OD_600_) were measured using a Tecan Infinite M200 PRO (Tecan Trading AG, Switzerland) at 0, 2, 4, 6, 8, 16 and 18 hours. At each time point, 200 μL of cell suspension was collected from each of three independent cultures and assayed on the counter. Cell concentration (with a 1:1 dilution of glycerol:sample) was calculated by multiplying the sum of the cell counts from ten images by a factor of 14,342 cells/mL as determined by the volume of the slide and the dilution factor from the sample preparation.

We used 18 bacterial isolates grown from a collection that had been isolated and characterized from the switchgrass (*Panicum virgatum*) rhizosphere (**Table 1**) (21). Each isolate was streaked from glycerol stock onto Reasoner’s 2A (R2A) agar + 0.5% succinate at 30 °C, and then single colonies were transferred to R2A broth at 30 °C with shaking at 200 rpm. Each suspension was diluted to an OD_600_ of 0.1 and counted as described above.

**Table 1.** Switchgrass rhizosphere bacterial isolates from the published collection of Grady and Shade 2024 (21) that were used in this study.

| Isolate ID | Genus | species | NCBI Accession | SRA Accession |
| --- | --- | --- | --- | --- |
| PvR115 | *Agrobacterium* | *rhizogenes* | OR754798 | SRR27348767 |
| PvR108 | *Chryseobacterium* | *lactis* | OR754795 | SRR27348770 |
| PvR079 | *Dyella* | *koreensis* | OR754774 | SRR27348793 |
| PvR102 | *Dyella* | *marensis* | OR754789 | SRR27348777 |
| PvR061 | *Enterobacter* | *ludwigii* | OR754766 | SRR27348812 |
| PvR147 | *Leclercia* | sp. | OR754808 | SRR27348798 |
| PvR062 | *Mycobacterium* | *hackensackense* | OR754767 | SRR27348811 |
| PvR096 | *Mycolicibacterium* | *mucogenicum* | OR754784 | SRR27348782 |
| PvR045 | *Novosphingobium* | *lindaniclasticum* | OR754756 | SRR27348823 |
| PvR090 | *Novosphingobium* | *capsulatum* | OR754779 | SRR27348788 |
| PvR052 | *Paenibacillus* | sp. | OR754758 | SRR27348821 |
| PvR098 | *Paenibacillus* | sp. | OR754786 | SRR27348780 |
| PvR021 | *Pantoea* | *agglomerans* | OR754742 | SRR27348838 |
| PvR083 | *Pseudomonas* | *frederiksbergensis* | OR754775 | SRR27348792 |
| PvR040 | *Rhodococcus* | *qingshengii* | OR754751 | SRR27348829 |
| PvR122 | *Roseococcus* | *suduntuyensis* | OR754803 | SRR27348803 |
| PvR112 | *Sphingobacterium* | *cladoniae* | OR754796 | SRR27348769 |
| PvR101 | *Sphingobium* | *indicum* | OR754788 | SRR27348778 |

### Soil assays

#### Cell separation from soil

To separate cells from the soil matrix, we modified a density gradient separation protocol based on previously published protocols (18, 22, 23). We disrupted 6 g of soil in 25 mL of 50 mM tetrasodium pyrophosphate 0.5% Tween80 (TSP-T) by blending (Waring, Eberbach Corporation, Michigan USA) it at high speed for 1 min. The slurry then was centrifuged 1000 × g for 10 min to pellet large soil particles (the centrifuging temperature depends on the soil original temperature) and to recover the supernatant. Supernatant from 1 gram of soil was slowly layered on top of 5 mL of 1.0 g/ml Nycodenz solution (Serumwerk Bernburg AG, Germany) and then centrifuged for 40 min at 12000 × g. We transferred 2 mL of the upper phase of Nycodenz containing microbial cells into a sterile 2 mL tube and centrifuged it at 12,000 × g for 10 min at 4 °C or the original temperature of soil samples. Then, a second extraction was performed by mixing 25 mL of TSP-T with the soil particles pelleted from the first extraction. The supernatants from two extractions were carefully discarded, and the pellet was resuspended in 1 mL of tetrasodium pyrophosphate (TSP, 50 mM). From each treatment, 10 μL of (diluted as needed) cell suspension was stained either with SYBR and RSG as described above. To calculate the concentration of cells per gram of dried soil, we multiplied the cell counts by the dilution factor and by the Quantom TX conversion factor (14319) and then divided by mass of the dry soil that used in the Nycodenz separation. Soil moisture content was expressed as the mass of water relative to the mass of fresh soil and determined by drying the soil overnight in an oven at 105°C.

To determine the recovery efficiency of the cell extraction from the soil matrix, we modified a protocol that was previously applied to plate-based assays of soil cell quantification (18). Briefly, we added a known concentration of the Green Fluorescent Protein (GFP)-tagged strain *Azospirillum baldaniorum* Sp245 P(ppdC-egfp) (24) to an alpine soil (site Galibier, coordinates: 45.0618, 6.4048, sandy-loam). *A. baldaniorum* was grown in Luria Bertani (LB) media overnight at 28 °C, and its concentration was measured using the QUANTOM Tx ^TM^ microbial cell counter. The culture was diluted with 1X PBS to 6.5×10^6^ cells/mL. One hundred (100) μL of the dilution was added to one gram of soil before the cell separation protocol. The recovery efficiency was calculated as recovered GFP-labeled cells divided by total added cells.

#### Lysozyme-resistant cell population

The fraction of lysozyme-resistant cells in the soil was quantified using a published protocol (25). Soil was collected from three locations in cold and warm seasons (**Table 2**), and the soils from the cold collection were used in this assay. After separating the cells from the soil using the Nycodenz protocol, 10 μL of the diluted cell suspension was stained separately with SYBR and RSG to obtain the features of the community (i.e., untreated). Next, 500 μL of diluted cell suspension was pelleted and re-suspended with 1.5 mL of fresh lysozyme buffer (1 mg/mL of lysozyme in a buffer: 50 mM Tris-HCl, 10 mM EDTA, and 1% Triton X-100), incubating at 37 °C for 100 min, with shaking at 100 rpm. Then, the lysozyme-treated samples were diluted and stained with SYBR or RSG to count the remaining lysozyme-resistant cells. This treatment was expected to enrich for spores and other hardy cells.

**Table 2.**
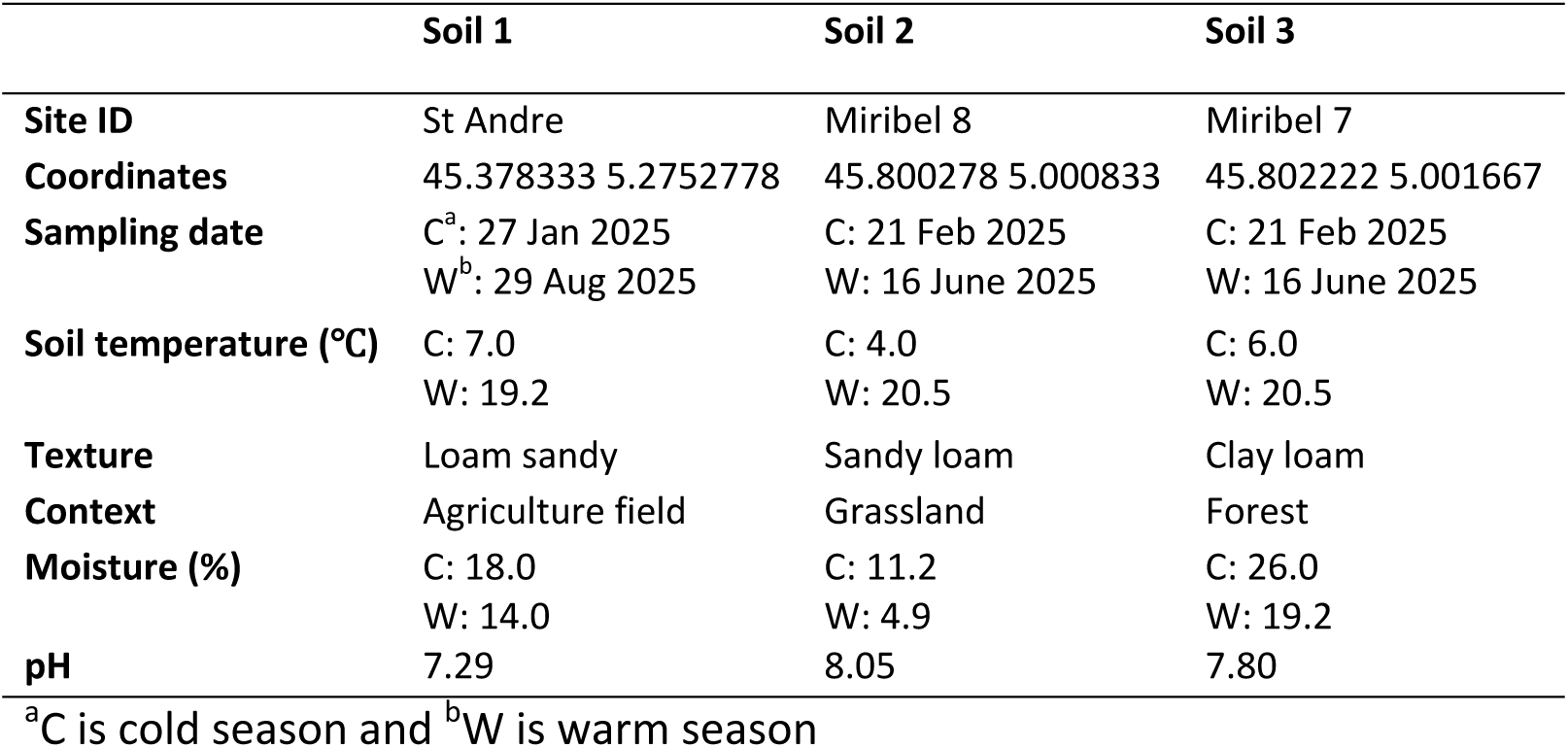
Metadata of the soil used in the respiration assays.

#### Relationship between cell redox activity and soil respiration

The relationship between soil community activity and respiration rate was assessed using French soils collected from three locations, each sampled in cold and warm seasons (**Table 2**). Fresh soils were stored after sampling at their original temperatures in temperature-controlled incubators in the dark until processing. Soils were sieved through 4 mm mesh and then incubated at 25 °C. Measurements were performed after 0, 15, and 21 days. Each soil sample included four biological replicates at each time point for a total of 48 samples.

Bulk soil respiration (ppm CO_2_) was measured using a gas chromatograph (Micro GC R3000, SRA Instrument, Marcy L’Etoile, France) as adapted from (26). Four replicates of 15.0 g of fresh soil were placed in a new glass plasma flask (150 mL) that was hermetically sealed with a rubber stopper. After a thirty-minute acclimation, automatic measurements were taken every 30 min for 7 h at 25 °C.

To estimate the rate of respiration from the evolution of CO_2_ concentration over time (the amount of carbon dioxide produced per gram of soil dry weight per hour), we assumed that an unknown concentration of limiting substrate (C_0_) at the beginning of the experiment and converted to CO_2_ at a constant rate (k) over time (t) by the soil population in the sample after a delay (d) when recording started. Therefore, we fitted the following first order kinetic model: CO_2_(t) ∼ C_0_ * (1 - e^(-k^ * ^(t + d))^) with a Gaussian sampling distribution. We also estimated the sum of the fluorescence signal from all the RSG stained cells in each soil using a generalized linear model with fixed effects for each soil, sampling season, and time after acclimatation, with a log-normal sampling distribution. Finally, we estimated the linear relationship between the estimates of the bulk respiration rate and substrate concentration against the estimates of the RSG fluorescence for each soil.

#### Soil heating experiment

We performed a proof-of-concept experiment with fresh soils to assess how bacterial community size and redox activity respond to heat stress, providing insights into ecological responses. Soils were collected on 19 August 2024 from a temperate site, Miribel Jonage Park (“Miribel 7”, coordinates: 45.8022, 5.0017, sandy clay loam) and at 22 August 2024 from an alpine site, Galibier (coordinates: 45.0618, 6.4048, sandy loam), both within the Rhône-Alps region, France. Soils were collected from 0-20 cm depth, sieved at 4 mm, and maintained at their original temperatures prior to conducting the assay (Miribel: 18 °C, Galibier: 12 °C). Six replicate mesocosms, each containing thirty grams of soil, were incubated at either their origin soil temperature or exposed to a heat treatment in a temperature-controlled and light-proof incubator (New Brunswick Innova 42R, Eppendorf). Both incubations were maintained at ambient soil moisture (Miribel = 16 %, Galibier = 20 %). For the heat treatment, the incubator temperature was cycled for 12 h at 35 °C and 12 h at 45°C for a total of 72 hours to simulate a diurnal cycle of a strong heatwave. Soil samples were collected after incubation and cells were immediately separated using the Nycodenz protocol and stained.

### Data analysis

All data processing, analyses, and figure preparation were done in the R environment (version 4.5.2) (27). Statistical analyses were done using custom generalized linear models and Bayesian sampling developed for each experiment using the *brms* (version 2.23.0) (28) and *RStan* (version 2.38) (29) packages. Bayesian sampling was done for each model using 16 independent Markov chain Monte Carlo collecting 1,000 samples each after warmup. Defaults *brms* priors were used unless noted otherwise. Model convergence and proper chain mixing were verified using the recommended diagnostics (https://mc-stan.org/docs/reference-manual/analysis.html).

### Data, Metadata and Code Availability

Raw images (.tif) and image-processed data with metadata (.csv) are available on FigShare (https://doi.org/10.6084/m9.figshare.30883721). The code for the image analysis is available on GitHub at https://github.com/DatInsights-com/process_quantom. The code for statistical analysis and R markdown notebooks for reproducing figures are available on GitHub at https://github.com/ShadeLab/MATRIX_CellQuantification_2026.

## Results

### Optical density is not a reliable metric to estimate bacterial population density and activity

First, we expected that cell staining with automated image analysis would recapitulate expected growth dynamics in pure culture. Specifically, total and active cells were expected to be highest and most similar in their values during exponential growth when cells are replicating, with total cells declining into stationary phase as well as the proportion of active cells. We sampled typical batch pure cultures of *Escherichia coli* and *Pseudomonas protegens* to measure optical density and enumerate total and active cell counts using our protocol. Replicates were highly reproducible for all the individual cell variables observed, enabling a well-powered and high-confidence analysis afforded by the large number of individual cells measured per replicate. Total cell counts (stained with SYBR) mirrored the increase in optical density as expected (**Figure 1A**), but the relationship between optical density and cell concentration is not linear and not identical between the two bacterial strains (**Figure 1B**). This demonstrates the limitations of using optical density to estimate cell concentrations. We obtained a reasonable fit between optical density and cell counts using a Gaussian sampling distribution and the following non-linear function for the mean: OD = OD_blank_ + (p1 * (total counts)^p2^), where p1 and p2 are fitted parameters (*E. coli*: p1 = 1.03E-2 [0.41E-2, 2.22E-2] 95% credible interval (hereafter “CrI”), p2 = 3.04E-1 [2.39E-1, 3.80E-1] 95%CrI; *P. protegens*: p1 = 9.97E-4 [2.60E-4, 31.0E-4] 95%CrI, p2 = 5.47E-1 [4.45E-1, 6.68E-1] 95% CrI).

**Figure 1.**
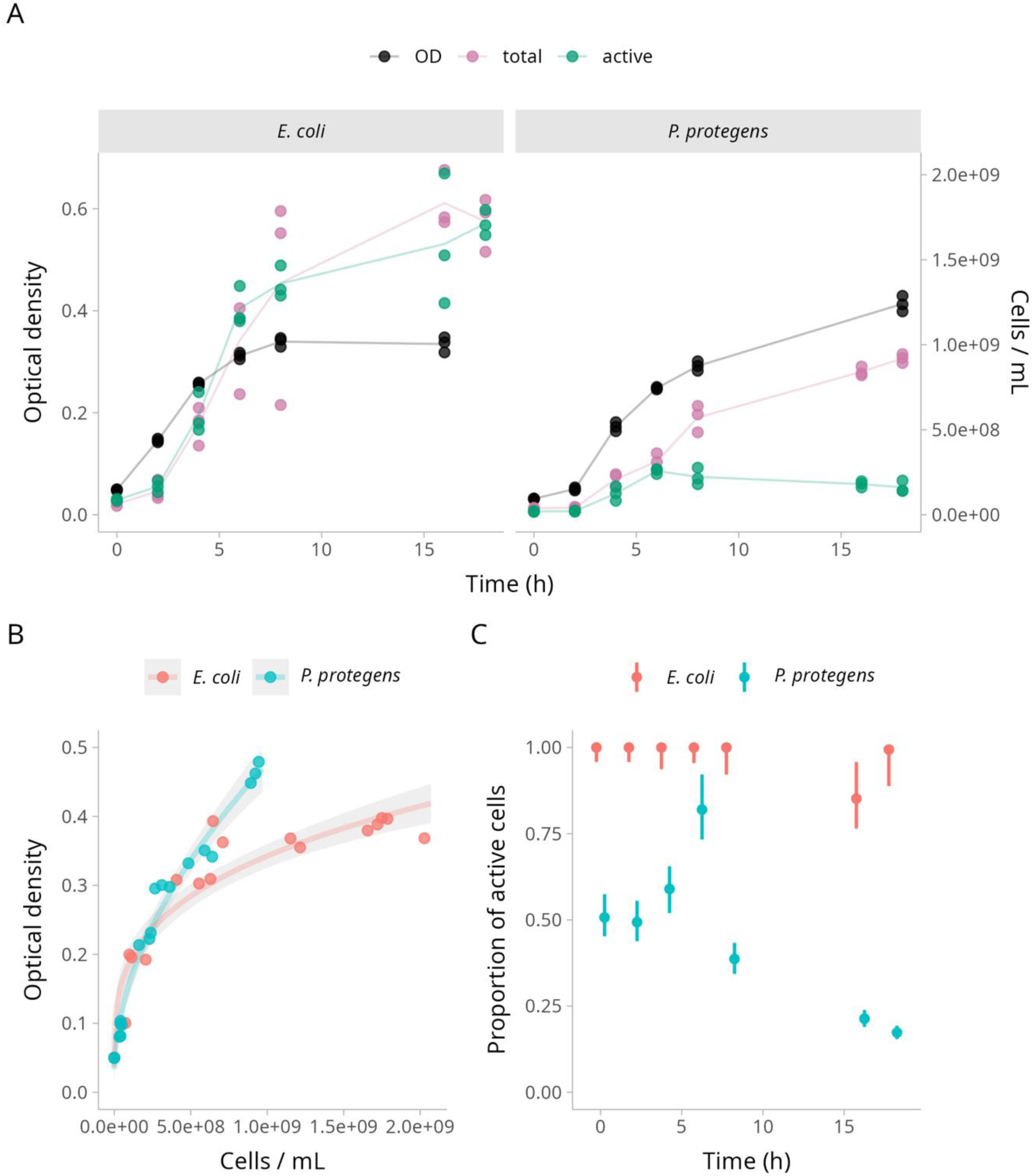
Quantification of total and active cell counts over time in growing bacterial batch cultures (n=3) of *E. coli* and *P. protegens.* A) Concentration of total (SYBR-stained, purple), respiration-active (RSG-stained, green) cells and optical density at 600 nm (OD_600_, black) over 18 hours. Each point represents an independent biological replicate. Optical density and counts are scaled on two different axes. Lines connect the means as a visual guide. B) Optical density as a function of total cell counts for each strain. The lines and gray areas represent the posterior probability distributions at the mode and 95% credible interval (CrI) of a fitted non-linear function. C) Proportion of active cells in each population for each time point inferred from the total and active cell counts. Points and lines represent the mode and 95% CrI of the posterior probability distributions for the estimated proportions.

The active cell counts (stained with RSG) revealed contrasting population dynamics between the two strains. For *P. protegens*, only a diminishing fraction of the population remained active after 5 hours of growth (**Figure 1A**). To estimate the proportion of active cells at each time point, we fitted a generalized linear model where total and active counts are modeled from a negative binomial sampling distribution with a mean parameter expressed as (total count) ∼ (mean total count) and (active count) ∼ (mean total count) * (proportion active). The posterior probabilities of the proportion of active cells reveals that close to 100% of *E. coli* cells are consistently active throughout the 18 hours of growth, whereas *P. protegens* starts around 50% of active cells, peaks at 70% at 6 hours and drops below 25% at 18 hours (**Figure 1C**). These results demonstrate that optical density or even total cell counts are not reliable metrics to estimate the functional size of growing bacterial populations.

In addition to cell counts, MATRIX provides quantitative information about the redox activity of the cells and morphological parameters. The distributions of fluorescence intensity for individual cells from the RSG stain shows (**Figure 2A**) a higher redox activity for *E. coli* at early time point, as expected when oxygen is readily available in the medium, but otherwise stable and unimodal distribution. However, there was an overall lower redox activity for *P. protegens* with a notable increase in variance, consistent with a population with heterogeneous activity. Morphological parameters appeared to be more correlated with the growth phases of the populations (**Figure 2B,C**). Both cell eccentricity and volume increase at early time points before decreasing in stationary growth. To quantify the correlation between cell volume and growth rate more precisely, we first estimated the culture growth rate from the total cell counts over time (**Figure 1**) by calculating the posterior probability distributions of the derivative of the change in cell numbers. We then fitted a linear model of the mean of the logarithm of the cell volume for both populations at each time point against the growth rate with a Gaussian sampling distribution. The results show the MATRIX protocol was able to capture the expected relationship between growth rate and cell size with very high confidence (slopes for *E. coli* and *P. protegens* = 19.0 [16.6, 21.9] 95% CrI) (**Figure 2D**).

**Figure 2.**
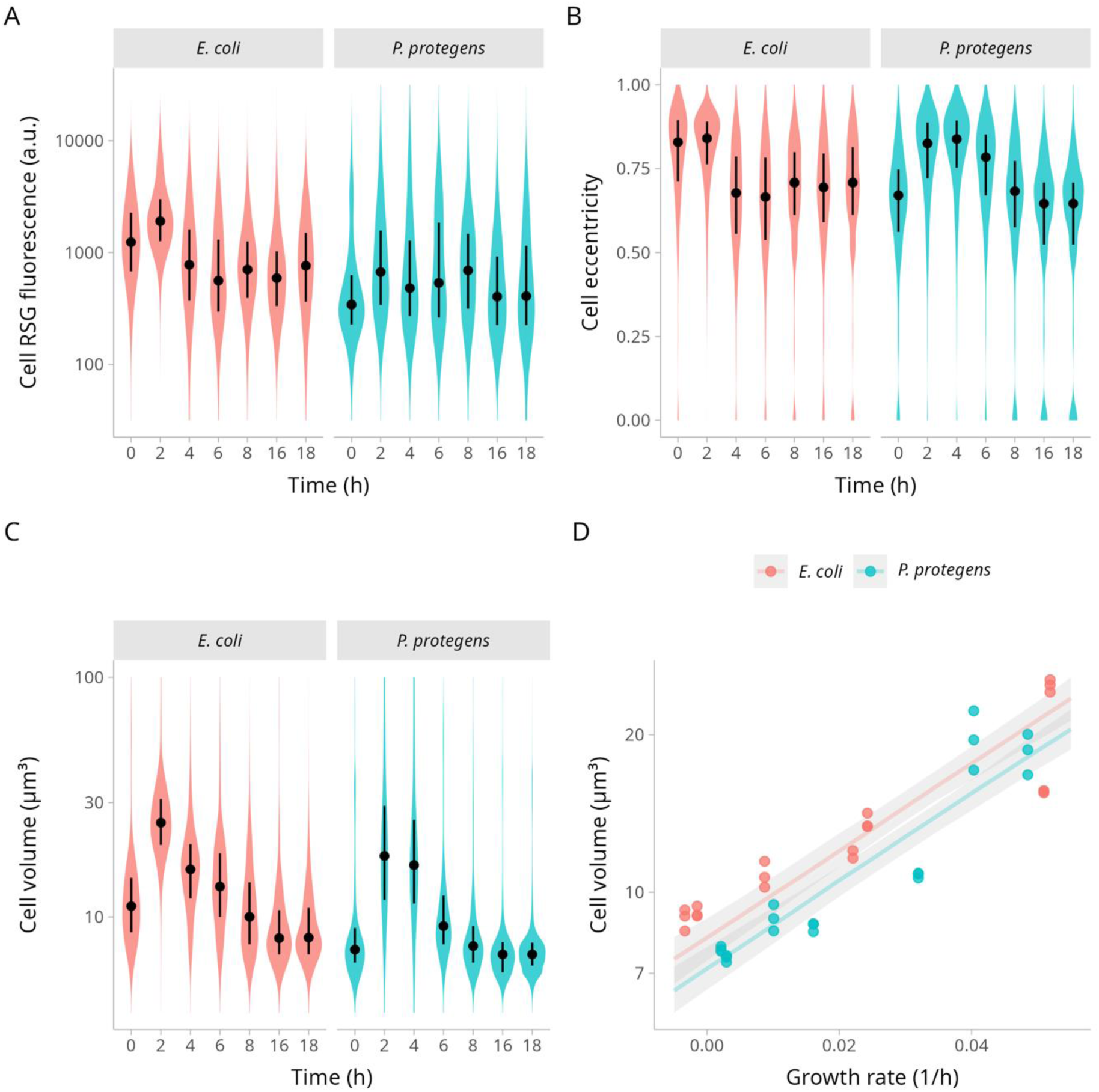
Distributions of single cell parameters from growing bacterial cultures (n=3) of *E. coli* and *P. protegens.* Each distribution represents the combined populations from the three biological replicates. A) Fluorescence intensity of RSG-stained individual cells over time, representing the total RSG activity of the cell. B) Cell eccentricity and C) volume over time. The black points and lines represent the medians and 50% quantiles of the populations. D) Average cell volume as a function of the estimated growth rates of each population. Volume is plotted on a logarithmic scale. The lines and gray areas represent the mode and 95% CrI of the posterior probability distributions of the linear fit.

### Counts and morphological parameters can vary wildly across environmental isolates

We next assessed the distributions of total and active cell counts for environmental bacteria that were isolated from the rhizosphere of switchgrass. Even though all cultures were diluted to the same equivalent optical density (0.1 at 600 nm) prior to measuring parameters (as sometimes used for assembling synthetic communities), they were not expected to have equivalent concentrations of cells due to inherent differences in growth dynamics and cell morphologies. As observed for *E.coli* and *P. protegens*, both the total cell counts and the proportion of active cells in a population vary significantly from one strain to another even at identical optical density (**Figure 3A,B**). The fluorescence intensity from the RSG-stained cells, as well as morphological parameters also varies significantly among populations (**Figure 3C,D,E**). These results illustrate how MATRIX can provide rich context to distinguish among different populations and inform practicalities for constructing synthetic community or co-culture experiments.

**Figure 3.**
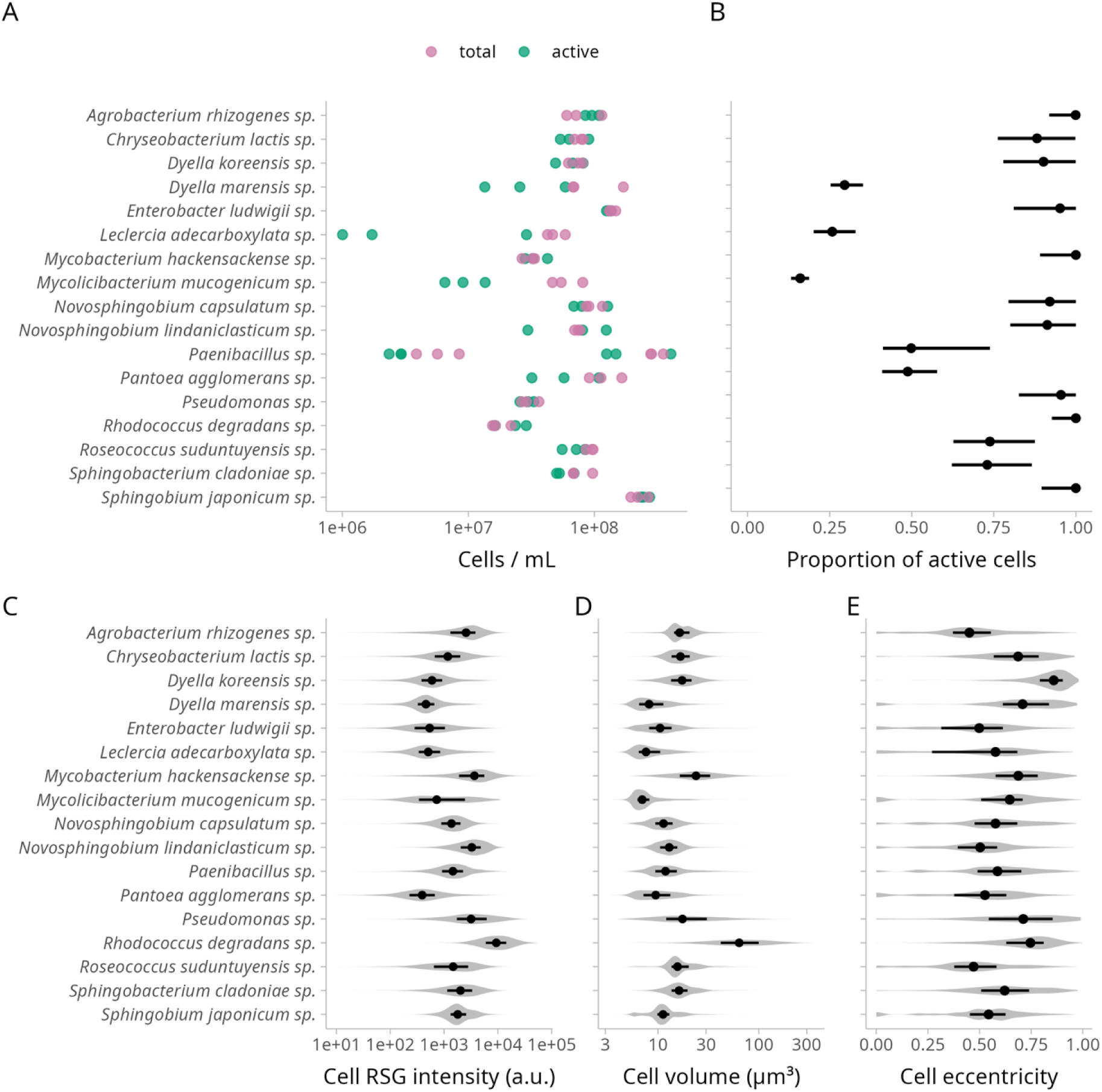
Comparison of total and active cell counts, redox activity, and cell morphology across 18 environmental isolates grown in triplicate and diluted to an optical density at 600 nm of 0.1. A) Concentration of total and active cells. B) Estimated proportion of active cells in each population. Points and lines represent the mode and 95% CrI of the posterior probability distributions. C) Distribution of fluorescence signal from RSG-stained cells across isolates. D) Distributions of cell volumes and E) eccentricity. The distributions combine the data from the three replicates. The points and lines represent the medians and 50% quantiles.

### Some but not all lysozyme resistant cells from soil show detectable redox activity

To test the efficiency of our protocol in recovering cells from soil samples we spiked soil with a known quantity of *A. baldaniorum* expressing the green fluorescence protein. After cell extraction from the soil matrix and counting we estimated the recovery rate to be 75% ± 18% (standard deviation) of cells. We observed no auto-fluorescence from the background soil debris.

We then evaluated the proportion of cells found in soil that are resistant to lysozyme treatment, which provides an indication of the proportion of spore and other hardy cells (25). Total and active cell counts were measured before and after lysozyme treatment in four independent replicates (**Figure 4**). The treatment decreased the total cell counts: Miribel 7 by 24% [12%, 36%] 95% CrI, Miribel 8 by 23% [11%, 34%] 95% CrI, St Andre by 21% [8%, 34%] 95% CrI (**Figure 4A**). The proportion of active cells had the following ranges: for Miribel 7 from 77% [66%, 90%] 95% CrI to 53% [40% 70%] 95% CrI; for Miribel 8 from 68% [58%, 80%] 95% CrI to 46% [36% 60%] 95% CrI; and for St Andre from 74% [62%, 86%] 95% CrI to 53% [41% 69%] 95% CrI, after treatment. Therefore, not all redox active cells were lysozyme sensitive, or not all lysozyme resistant cells were inactive. We estimated the proportions of each of the four categories in the original soil samples: resistant and inactive (RI), resistant and active (RA), sensitive and active (SA), and sensitive and inactive (SI) (**Figure 4B**). With the constraint that no proportion can be negative we found that SI = 0% in all three soils. These results indicate, first, that most lysozyme-sensitive cells were also active (e.g., there were no detected inactive and “vegetative” cells). Second, these data suggest that a large proportion of lysozyme-resistant cells were inactive. Furthermore, many lysozyme-resistant cells had detectable RSG signal and were classified as active. In fact, most detected active cells belonged to the resistant pool. Together, these results suggest that considering the activity level, rather than assigning a binary definition of active or not, could prove informative for understanding the active components of the soil microbiome.

**Figure 4.**
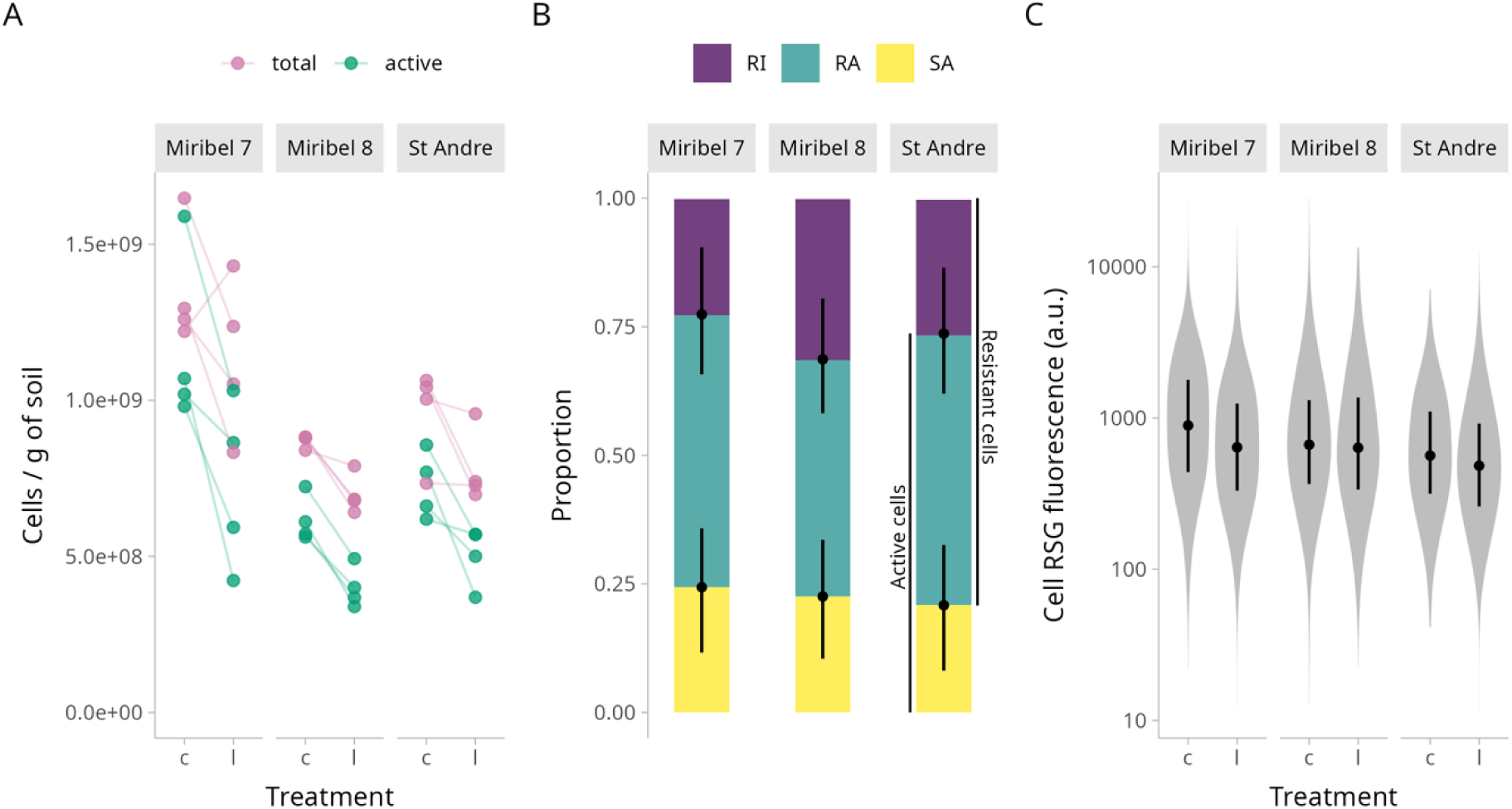
Effect of lysozyme treatment on total and active cell counts in a soil community. A) Cell concentration of untreated (c) and lysozyme-treated (l) soil sample. Each connected pair of points represents an independent biological replicate. B) Estimation of the proportions of cells from three category: resistant and inactive (RI), resistant and active (RA), and sensitive and active (SA). The points and lines represent the median and 95% CrI of the posterior probability distributions of the estimates. Active cells = RA + SA, resistant cells = RA + RI. C) Distributions of cell RSG fluorescence signal across soil samples and treatments. The points and lines represent the medians and 50% quantiles of the distributions.

### Total fluorescence from RSG stained cells predicts bulk respiration rates

To test if our protocol can predict the bulk rate of soil respiration activity, a commonly measured ecosystem function, we measured the respiration rates of three soils collected from various geographic locations each during a cold season (soil temperatures between 4 and 7 °C) and a warm season (temperatures between 19 and 21°C) , and compared them to the fluorescence signal of RSG stained cells isolated from the same soils. We found a positive relationship between the total RSG fluorescent signal and both the rate of respiration (slope = 6.20E-2 [2.87E-2, 9.43E-2] 95% CrI) and the amount of substrate available (slope = 1.25E-1 [0.62E-1, 1.92E-1] 95% CrI) (**Figure 5**). These results suggest that the RSG signal per soil are related to and may indicate soil bulk respiration.

**Figure 5.**
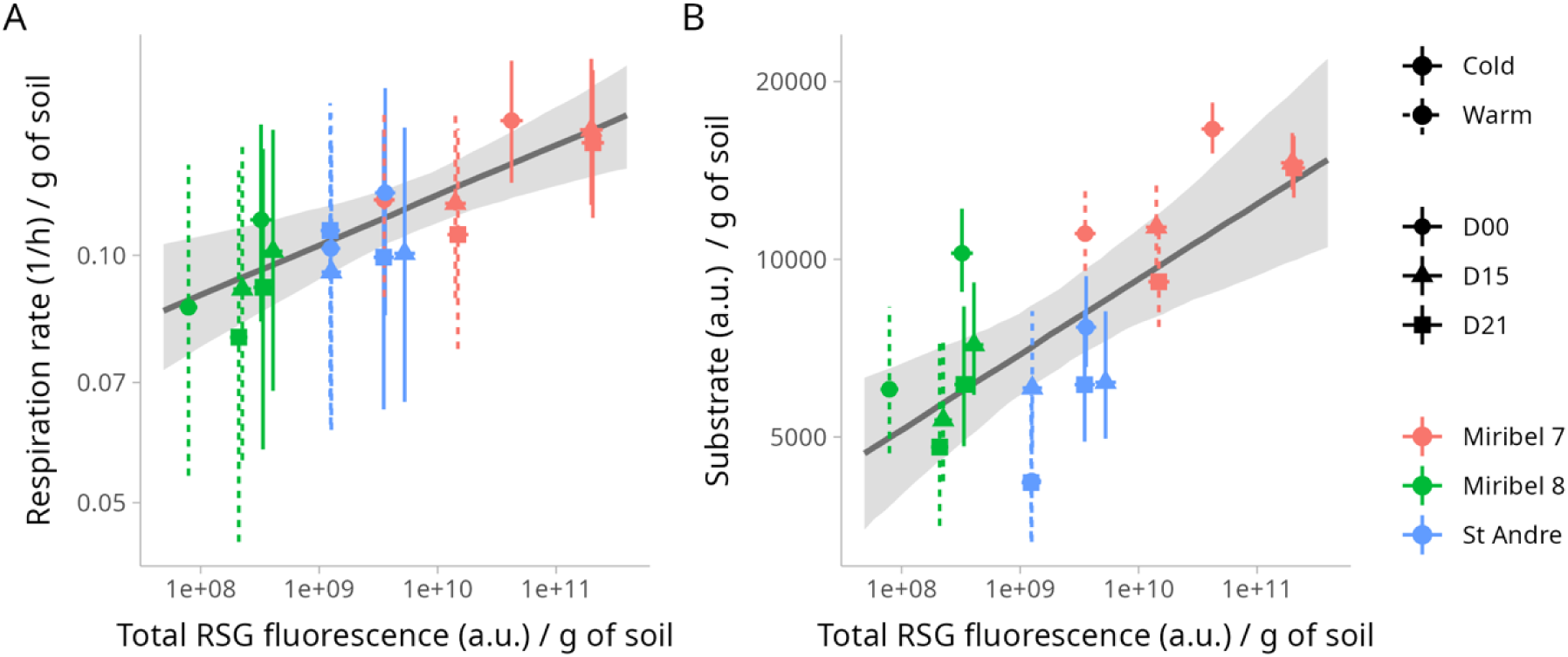
Relationship between total RSG fluorescence signal from cells extracted from soil samples and bulk respiration parameters. A) Bulk respiration rate as a function of total RSG fluorescence. B) Initial substrate concentration as a function of total RSG fluorescence. The points and lines represent the medians and the 95% CrI of the posterior probability distributions for each estimate. Each color represents a sampling location, shapes represent the duration of soil acclimatation, and line style distinguishes the season for soil sampling. The regression line and gray area are the median and 95% CrI of the posterior probability distribution of the linear fit. The data is plotted on a double logarithmic scale.

### Heat treatment

To test the soil population response to heat perturbation using MATRIX, we measured the total and active cell counts from two soils sampled at different locations after exposure to high heat stress. Heat treatment increased the total concentration of cells in soil from Galibier (1.9X) but not Miribel (1.0X) (**Figure 6A**). The proportion of active cells was between 50% and 60% on average, but heat stress increased the proportion above 60% in the Galibier sample and above 80% in the Miribel sample (**Figure 6B**). On the other hand, the distributions of fluorescence signal from RSG stained cells were not different between the samples or affected by heat stress (**Figure 6C**). Comparatively, the Galibier soil had a relatively larger increase in total counts but a modest increase in the proportion of active cells, while Miribel had more stable total cell counts but a larger increase in the proportion of active cells after heating. Both soils shared equivalent ranges of activity intensity, suggesting that either response ultimately supported similar activity levels. These results illustrate that accounting for both active and total cells can provide specific insights into different soil responses that may otherwise be masked without consideration of both components with activity intensity.

**Figure 6.**
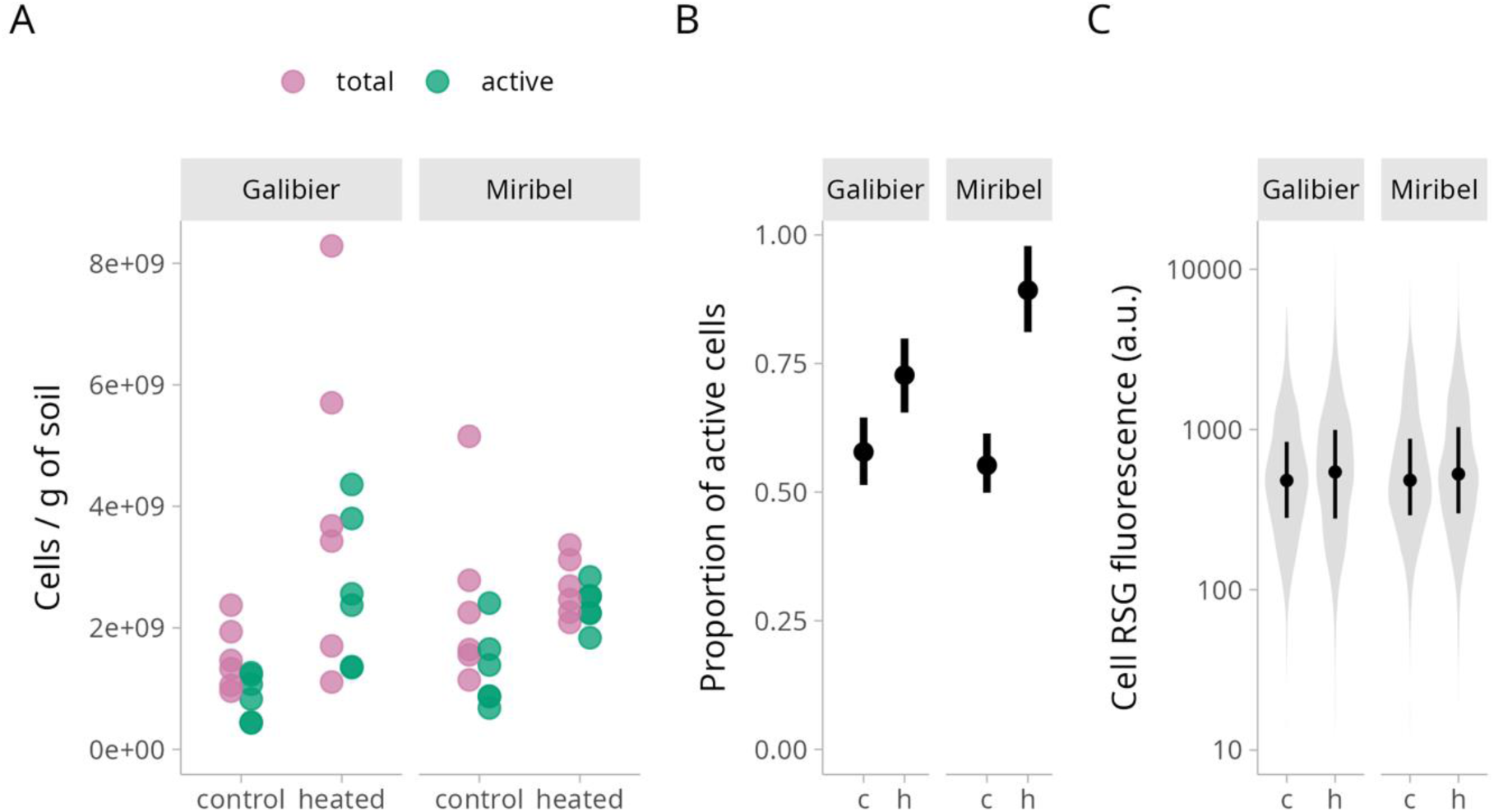
The impact of high temperature stress on microbial communities from two soil samples. A) Total (purple) and active (green) cell counts per gram of dry soil. Each point represents an independent biological replicate. B) Estimated proportion of active cells in each sample. Points and lines represent the mode and 95% CrI of the posterior probability distributions. C) Distributions of fluorescence signal from individual cells stained with RSG. Points and lines represent the median and 50% quantiles of the distributions.

## Discussion

MATRIX provides rapid, direct quantification of total and active cells and yields phenotypic information that strengthens ecological interpretation of microbiome datasets. Our validation across pure cultures, environmental isolates, and soil communities demonstrates that MATRIX reliably captures expected growth dynamics, predicts bulk respiration, and detects treatment-induced shifts in community size and activity. Together, these findings show that the workflow recovers biologically expected patterns while providing phenotypic resolution not accessible with OD-based or DNA-based methods. Overall, MATRIX has the potential to enrich environmental microbiome experiments by enabling direct quantification of cell, population, and community data, information that is indispensable for achieving robust ecological interpretation of microbiome responses. Direct measurements of population and community size anchor microbiome analyses in fundamental ecological accounting (1).

MATRIX offers several advantages. First, it allows rapid, direct quantification of total cell counts within a microbiome sample, avoiding the biases of cultivation (e.g., “the great plate count anomaly”, (30)(31)), and of indirect molecular approaches (e.g., sampling, cell lysis, nucleic acid extraction, molecular protocol biases). It also provides an independent, quantitative baseline against which ‘omics data derived from the same sample can be standardized, facilitating a shift from purely compositional and to more quantitative microbiome profiling. Beyond this, MATRIX generates rich, single-cell phenotypic profiles that can provide insights into direct cellular responses to treatment (e.g., changes in size, volume, eccentricity). When used with RSG staining, the workflow also enables assessment of changes in the active fraction of the community, which is important in perturbation experiments and similar studies that are expected to result in substantial shifts in activation and in death, including experiments investigating how microbial functionality may change climate disturbance (16, 32).

When comparing proportions of active cells across conditions, simply pooling all cells and using standard tests can lead to numerical issues and inaccurate estimates, because total counts and active counts are measured from two independently stained subsamples derived from a shared biological sample. To accommodate this nestedness and the inherent correlation between total and active counts, we developed a Bayesian hierarchical model that estimates total counts and the proportion of active cells jointly, rather than treating SYBR and RSG counts independently. This approach provides major advantages because it enables partial pooling, properly accounts for intra-group correlations, handles unbalanced designs without artificial data manipulation, and naturally constrains proportions between 0 and 1, regardless of random effects during sampling. As a result, it yields more accurate and reliable estimates that reflect the experimental structure and variance.

There are also considerations associated with each step of the protocol. Density-gradient separation for cell extraction efficiently removes most soil particles (18, 33, 34), but, like all physical separation methods, it does not yield complete cell recovery (35) and can introduce biases. Reported extraction efficiencies vary across soils (33)(18), and our two-step recovery of ∼75% falls within the expected range of 55-80% recovery (33). A third extraction did not significantly improve yields (data not shown), suggesting diminishing returns. Environmental samples with differing recovery efficiencies may incorporate include these uncertainties into statistical models for direct comparisons. Cell extraction is also influenced by cell morphology and density, potentially biasing against organisms with low-density cell structures or mycelial forms (36). Similarly, particles with densities similar to bacteria, such as free chloroplasts or mitochondria, may be extracted; although we did not observe cell autofluorescence suggestive of chloroplasts in the bulk soil, chloroplast filtering from the data (based on size and shape) may be necessary for plant-associated microbiomes studies.

As with other fluorescence-based protocols, SYBR and RSG staining exhibit known biases in nucleic acid visualization and redox-activity detection (17). Although prior work shows that RSG does not alter cell physiology (19, 37), its signal still reflects both biological variation and methodological sensitivity. The method also has a minimum detectable cell size imposed by image resolution, making it unsuitable for very small cells such as nanobacteria.

A consideration specific to RSG is the need to minimize the processing timing before measurement, or to use a fixative, to preserve the activity signal so that the active fraction reflects the most immediate environmental state, which is not directly knowable. Because reductases and metabolic enzymes decline gradually after cell death (38), RSG can detect recently active cells for a window of a few hours (39–41). Given the density-gradient extraction prior to staining, assayed cells likely have minimally compromised membranes, limiting signal from long-dead cells and removing free DNA.

Instrument and software considerations also apply. While the QUANTOM Tx counter likely performs well for homogenous clinical cultures, we found that its built-in software was less precise for mixed growth stages or environmental communities with diverse phenotypes. We therefore developed the MATRIX imaging analysis protocol to provide transparent quality control and tunable parameters tailored to complex samples. This flexibility improves detection but introduces additional analytical steps that require user validation.

The laboratory protocol (extraction, staining, and imaging) can be completed in about two hours, and up to eight samples can be processed in parallel. At the time of writing, the cost of the assay was about six Euros (approximately seven USD) per sample for both total and active counts. Future optimizations may expand the method to detect and differentiate other microbes like archaea and fungi, account explicitly for plastids, and improve identification of more irregular microbial morphologies.

We strongly advocate that active and total community sizes be routinely measured and reported for environmental microbiome studies, whether by MATRIX or other workflows. Doing so will improve understanding of the size and variability of the dormant pool (“seed bank”(15)) across space, time, and experimental conditions, which is fundamental information for evaluating its potential to support microbiome resilience (42).

MATRIX provides a quantitative foundation for microbiome studies, enabling direct assessment of population and community size and improving interpretation of ecological and functional responses. Extending these tools across hosts and ecosystems will support more mechanistic, activity-informed microbiome and microbial systems science.

## Acknowledgements

This research was funded by by the European Union (ERC, MicroRescue, 101087042). Views and opinions expressed are however those of the author(s) only and do not necessarily reflect those of the European Union or the European Research Council. Neither the European Union nor the granting authority can be held responsible for them. We also acknowledge the Agriculture and Food Research Initiative (AFRI) of the United States Department of Agriculture (USDA) National Institute of Food and Agriculture (NIFA) Grant number #2024-67019-42477, which supported the assays involving the switchgrass rhizosphere isolates. We thank Claire Prigent-Combaret for sharing the GFP-labeled *Azospirillum baldaniorum* strain, Miribel Jonage Park for access for soil collection, and Juliana Almario and Lauren Gillespie for access and help with soil collection for the Alpine Galibier soil associated to the Zones Atelier Alpes. Finally, we acknowledge our unit’s Microbial Activities in the Environment (AME) research support platform and Jonathan Gervaix for soil respiration measurements.

